# Cryo-EM structure of MutL-activated MutH

**DOI:** 10.64898/2026.04.21.719898

**Authors:** Emma Arean Ulloa, Meindert Hugo Lamers

## Abstract

Strand discrimination after mismatch recognition is the second essential step that is unique to DNA mismatch repair. MutH is a latent endonuclease that cuts the newly synthesized strand at hemi-methylated GATC sites upon activation by MutL, which acts as a signaling protein between mismatch recognition by MutS and strand discrimination by MutH. While much is known about the recognition of mismatches by MutS, little is known about the activation of strand incision by MutH. Here we present the cryo-EM structure of MutL bound to MutH on a hemi-methylated GATC site at 2.8 Å resolution. MutL binds MutH at the back of the protein, 25 Å away from the active site. The structure reveals that activation occurs through an allosteric pathway that acts through a push-and-pull mechanism that releases an inhibitory loop from the DNA minor groove enabling histidine 112 to pull the DNA backbone closer into the active site where a lysine 116 positions a water molecule for the nucleophilic attack at the scissile bond 5’ to the GATC sequence. Thus, our structure reveals how MutL couples mismatch recognition by MutS to daughter strand discrimination by MutH.

## INTRODUCTION

The faithful replication of genetic information requires the correction of mispaired bases that occur during DNA synthesis. DNA mismatch repair (MMR) is a post-replicative pathway that recognises and corrects these replication errors, thereby reducing the overall error-rate 100-1000 fold (1). Defects in MMR lead to elevated mutation rates, contributing to the emergence of antibiotic resistance in bacteria and a predisposition to cancer in humans, including hereditary non-polyposis colorectal cancer (Lynch syndrome) and various sporadic cancers (2).

Unlike other DNA repair systems that recognize damaged DNA, MMR is unique in that it does not work on damaged DNA but recognizes bases that wrongly paired. Therefore, in addition to recognizing the mismatch, MMR also needs to discern between the template strand and the newly synthesized strand. In *Escherichia coli*, these first steps of MMR proceed through a coordinated cascade of protein–DNA interactions involving MutS, MutL, and MutH. As a first step in the process, MutS recognises the mismatched base pairs. Then, it undergoes an ATP-driven conformational changes that convert MutS in a sliding clamp which enables the recruitment of MutL to the DNA. MutL then transmits the activation signal to the latent endonuclease MutH that cuts the newly synthesized strand 5’ to hemi-methylated GATC site, directed by the methyl group on the template strand and the absence of a methyl group on the new strand. Following incision, the removal of the new strand and the mismatch contained therein is performed by the helicase UvrD and a 3’-5’ or 5’-3’ exonuclease (3–5).

While much is known about the initial step of mismatch recognition by MutS and recruitment of MutL, less is known about activation of the incision of the newly synthesized strand by MutH. MutH belongs to the superfamily of PD-(D/E)XK-fold nucleases that include type-II restriction enzymes, and various nucleases involved in DNA repair, DNA replication, and tRNA intron splicing (6). In this large group of enzymes MutH is unique in that it selectively cleaves the unmethylated strand 5′ to the GATC site, but only when activated by MutL. The crystal structure of *Haemophilus influenzae* MutH bound to a hemi-methylated GATC DNA substrate (7) revealed how it selectively cuts the non-methylated strand but gives little clues on its activation by MutL. Biochemical and crosslinking studies indicate that the ATP-induced dimerization of the N-terminal MutL ATPase domains (MutL^LN40^) is required to bind and activate MutH (8, 9). In contrast, the role of the C-terminal dimerization domain of MutL (MutL^LC20^) is less clear (8–10). Importantly, how the interaction with MutL triggers the latent endonuclease activity of MutH is not understood (8, 11, 12).

In this work, we present the cryo-EM structure of the MutH-MutL complex bound to a 14 base pair hemi-methylated DNA duplex. Even though full length MutL was used, only the MutL^LN40^ dimer is visible in the complex, indicating that the MutL^LC20^ domains are not required for binding and activation of MutH. In our structure, MutH is positioned in a groove formed by the two MutL^LN40^ domains. Remarkably, the MutL^LN40^ dimer contacts MutH at the back of the molecule, > 25 Å away from the MutH active site. Comparison of our MutL-bound MutH-DNA complex to that of the MutL-free MutH-DNA complex suggests an allosteric activation mechanism in which the binding of the MutL^LN40^ dimer induces the displacement of an inhibitory loop in MutH from its position in the minor groove of the DNA, enabling the DNA to move closer into the active site, promoted by a hydrogen bond between histidine 112 and the DNA backbone. In this pre-incision state, lysine 116 coordinates a water molecule in a perfect position for an inline attack of the scissile bond 5’ to the GATC sequence of the non-methylated strand. This activation mechanisms bears similarities to the structurally related Type-II restriction endonucleases Sau3AI and EcoRII in which activation is achieved by dimerization, but in these instances by two identical monomers. Furthermore, the release of an inhibitory element from the minor groove of the DNA is also observed in other nucleases, indication that the activation of MutH by MutL is a variation of theme more generally used by type II restriction endonucleases.

## MATERIAL AND METHODS

### Protein purification

All proteins were expressed in *E. coli* BL21(DE3) cells grown in 2x YT media (Formedium, UK) at 30 °C. MutL was expressed for 2 hours and MutH for 3 hours. Cells were harvested and lysed by sonication and cleared by centrifugation at 24.000 x g for 40 minutes. MutL^WT^ and MutL^LN40^ (residues 1 – 331) were lysed in 25 mM HEPES pH 7.5, 500 mM NaCl, 10 mM imidazole, 5 mM MgCl_2_, and 1 mM DTT. The cleared lysate was loaded onto a HisTrap column (Cytiva) and eluted with a gradient of imidazole to 500 mM. Pooled fractions were diluted to 100 mM NaCl and further purified on a HiTrap Q column (Cytiva) using a NaCl gradient up to 1 M. A final purification step was performed using a Superdex 200 16/600 size exclusion chromatography column (Cytiva), equilibrated in 25 mM HEPES pH 7.5, 150 mM NaCl, 5 mM MgCl_2_, 2 mM DTT. Purified proteins were snap-frozen in liquid nitrogen and stored at −70 °C. Cells expressing MutL^LC20^ (residues 350-615) were lysed in 25 mM Tris pH 8.0, 500 mM NaCl, 20 mM imidazole and the cleared lysate loaded onto a HisTrap column and eluted with a gradient of imidazole to 500 mM. Eluted fractions were diluted to 100 mM NaCl and applied to a Q column and eluted with a gradient to 1 M NaCl. The eluted protein was diluted to 100 mM NaCl and loaded onto a Heparin column (Cytiva) for final purification. Cells expressing MutH were resuspended in 25 mM Tris pH 8.0, 500 mM NaCl, 20 mM imidazole. Clarified lysate was loaded onto a HisTrap column and eluted with an imidazole gradient up to 500 mM. Pooled fractions were diluted to 100 mM NaCl and applied to a Q column and eluted with a gradient to 1 M NaCl. Following this, fractions containing MutH were pH-exchanged to pH 5.5 and loaded onto an Hitrap SP column (Cytiva) column and eluted with a gradient to 1 M NaCl. Final purification was performed by size-exclusion chromatography using a Superdex 75 column (Cytiva). All purified proteins were snap-frozen in liquid nitrogen and stored at −70 °C .

### Cryo-EM sample preparation and data acquisition

Purified proteins were diluted in 25 mM HEPES pH 7.5, 150 mM potassium glutamate, 5 mM MgCl_2_, 0.01% (w/v) Tween 20, 2 mM DTT and 3 mM AMPPNP. The complex was assembled by incubating 4 µM MutH^E56A^, 8 µM MutL (monomer concentration), 20 µM 14 bp DNA substrate (IDT) containing a methylated GATC sequence in one strand (5’-GCCAGGA^Me^TCCAAGG, where Me denotes the methylated adenosine), and a regular GATC sequence in the opposite strand (5’-CCTTGGATCCTGGC). Samples were incubated for 10 min on ice and then 4 µL of the sample was applied to glow-discharged Quantifoil R 2/1 300 mesh copper grids. Grids were glow discharged for 30 seconds at 25 mA using a PELCO easiGlow™ machine. The grids were blotted for 2 seconds at 98% humidity and 20 °C, then flash-frozen in liquid ethane using a Leica EM GP plunge freezer.

Cryo-EM movies were collected on a Titan Krios (FEI) electron microscope operating at 300 kV equipped with a Gatan K3 direct electron detector and a BioQuantum energy filter set to 20 eV. Images were recorded using EPU software (https://www.fei.com/software/epu-automated-singleparticles-software-for-life-sciences/) in counting mode. Details on electron dose, magnification, and effective pixel size are provided in Supplementary Table 1.

### Cryo-EM data processing and model building

All image processing was performed using cryoSPARC v4.7.0 (13) (Supplemental Fig. S2, Supplemental Table 1). Motion correction was performed using cryoSPARC’s Patch Motion Correction algorithm, and CTF parameters were estimated using the Patch CTF Estimation module. Initial particle picking was done using the blob picker, specifying a particle size range of 65–120 Å. Particles with normalized cross-correlation scores below 0.4 were discarded. The remaining particles were extracted using a box size of 256 × 256 pixels at 0.836 Å /pixel and downsampled to 96 × 96 pixels (2.224 Å/pixel) for faster processing. After 2D classification, well-defined classes were used to generate templates for a second round of particle picking using the template picker algorithm. A curated set of 565.026 particles from 2D classification were then used for ab initio 3D reconstruction. Following several rounds of heterogeneous refinement, curated particles were re-extracted with the full box size (256 × 256 pixels, 0.836 Å /pixel) and subjected to non-uniform refinement to produce a high-resolution reconstruction (14). The final particle set underwent reference-based motion correction and CTF refinement.

The final refinement was performed using non-uniform refinement, yielding a post-processed map sharpened by applying a B-factor of 150 to correct for the modulation transfer function of the detector. The final map achieved an average resolution of 2.83 Å, as determined by Fourier shell correlation (FSC) analysis. The final EM map included 565,000 particles.

Model building was performed using Coot (15) and refined in Phenix (16). In brief, model building began with rigid-body fitting of a composite PDB consisting of the known *E. coli* MutL^LN40^ crystal structure (PDB ID: 1B63) (17) and a model based on the structure of *Haemophilus influenzae* MutH (PDB ID: 2AOR) (7) into the experimental density map using Coot. Multiple rounds of refinement were performed in Phenix, with manual adjustments in Coot between iterations. The final model was refined and validated using Molprobity (18).

### MutH strand-incision assays

MutH endonuclease activity was assessed by denaturing gel electrophoresis. Reactions were performed at room temperature in buffer containing 25 mM Tris pH 7.5, 150 mM KCl, 5 mM MgCl_2_, 10% glycerol, 0.1 mg/mL BSA, 0.005% (w/v) Tween 20 and 3 mM ATP or AMPPNP. MutH, MutL, MutL^LN40^ or MutL^LC20^ were diluted to the indicated concentrations and incubated with a regular or hemi-methylated 14 bp DNA substrate containing the GATC recognition sequence (top strand: 5’-GCCAGGA^Me^TCCAAGG, where Me denotes the methylated adenosine when present, and a fluorescently labelled bottom strand: 5’-FAM-CCTTGGATCCTGGC) at 50 nM to initiate the reaction. At time points 0, 5, 10, 30, and 60 minutes, 5 μL aliquots were withdrawn and quenched in 95% formamide, 10 mM EDTA, 0.04% (w/v) xylene cyanol, followed by heating at 95 °C for 5 minutes. Reaction products were separated on a 18% (19:1) denaturing polyacrylamide gel containing 7.5 M urea, run in 1x Tris-Borate-EDTA buffer. The gel was pre-run at 30 W for at least 1 hour, followed by electrophoresis at 25 W for 30 minutes. Gels were scanned at 488 nm using a Typhoon™ scanner to visualize the FAM labelled DNA products.

### Size exclusion chromatography analysis

Oligomerization of MutL^FL^, MutL^LN40^, and MutH was analysed by size exclusion chromatography using a 2.4 ml (3.2/300) Superdex 200 Increase column (Cytiva). All runs were performed in 25 mM HEPES pH 7.5, 150 mM KCl, 5 mM MgCl_2_ and 1 mM DTT. For the AMPPNP induced dimerization of the LN40 ATPase domains, 3 mM AMPPNP was added to 4 µM of MutL^FL^ or 7 µM MutL^LN40^ and incubated for 10 or 85 minutes. For binding of MutL to MutH, 22 µM MutL (dimer concentration) and 22 µM MutH (monomer concentration) were mixed and incubated for 15 minutes on ice prior to injection. In all cases, 100 µL sample was injected into the column and resolved at 0.1 ml/min at 4 °C. Selected fractions were analysed by SDS-page on a 15% acrylamide gel.

## RESULTS

### Cryo-EM structure of the MutL-MutH complex on a 14 base pair hemi-methylated DNA subtrate

We determined the cryo-EM structure of the MutH–MutL complex bound to a 14-bp DNA duplex containing a hemi-methylated GATC site to 2.8 Å resolution (Fig. 1A-B). To prevent nicking of the DNA, we used a catalytically dead version of MutH in which glutamate 56 was replaced by an alanine (19), while the non-hydrolysable ATP analog AMPPNP was used to induce the active, closed form of MutL (9). The structure shows a MutL dimer and one MutH bound to DNA. A second, weaker density is observed on the opposite side of the MutL dimer (Fig. 1A, light purple), where a second MutH-DNA could bind. However, the density is weak and covers less than half of a MutH molecule. Attempts to separate particles with two MutH molecules and generate a separate 3D volume were not successful, indicating that the fraction of particles with two MutH molecules is too small to generate an independent 3D class.

**Figure 1.**
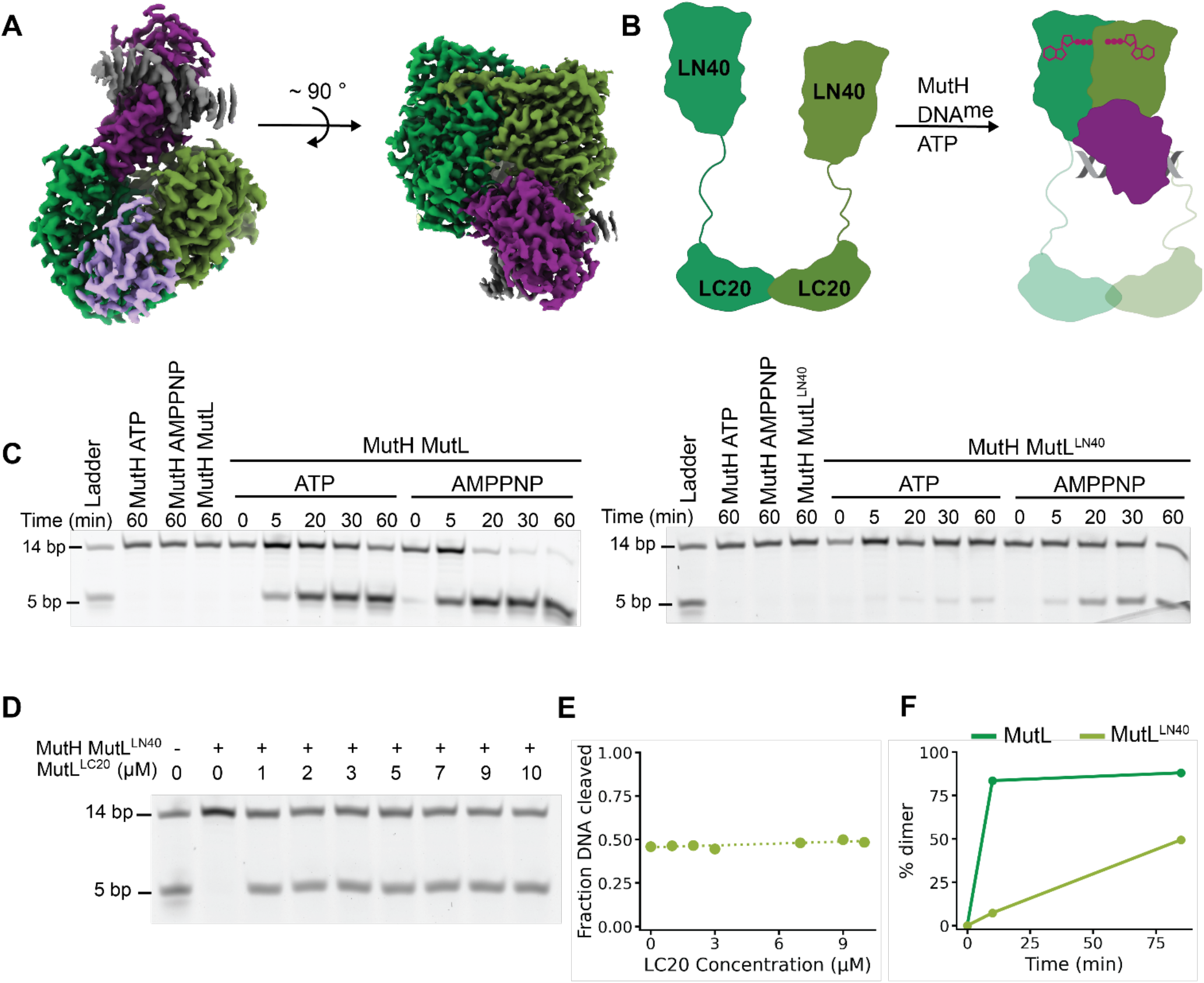
Cryo-EM structure of the MutL-MutH-DNA complex. **(A)** Top (left) and front (right) view of the MutL-MutH complex bound to a hemi-methylated DNA substrate. The MutL dimer (LN40 domains only) is shown in two shades of green, MutH is shown in purple, and the DNA in dark grey. The partial density for a second MutH monomer is shown in light purple (see main text for details). **(B)** Schematic representation of the formation of the MutL-MutH-DNA complex. Left: full-length MutL forms an obligate dimer via the C-terminal LC20 domains, while the N-terminal ATPase domain (LN40) do not interact in the absence of ATP. In the presence of ATP the LN40 domains dimerize and form a complex with MutH and DNA. The LC20 domains that are unresolved in the cryo-EM structure are depicted in transparent colour and dashed lines. ATP molecules are shown in red. **(C)** MutH DNA nicking assay showing the stimulative eOect of MutL^FL^ and MutL^LN40^ on the activity of MutH in the presence of ATP or AMPPNP. **(D)** Titration of LC20 domain of MutL (MutL^LC20^) to the MutH-MutL^LN40^ reaction. **(E)** Quantification of the nicked product shown in panel D at increasing concentrations of the LC20 domain. **(F)** Quantification of LN40 domain dimer population of MutL^FL^ and MutL^LN40^ at 10 and 85 minutes after incubation with AMPPNP as determined by size exclusion chromatography (see Supplementary Fig. S3).

Although full-length MutL was used during sample preparation, only the dimer of MutL^LN40^ was resolved in the cryo-EM map. The C-terminal MutL^LC20^ is separated by a long flexible linker (20), which could explain the lack of density for this domain in the cryo-EM map. The absence of MutL^LC20^ in the cryo-EM map suggest that this domain is not strictly required for activation of MutH. Indeed, previous work has shown that MutL^LN40^ is enough to activate MutH *in vitro* (8), yet other lines of work indicate that MutL^LC20^ may also play a role in the activation of MutH (9, 12, 20). Therefore, to investigate the role of MutL^LC20^, we performed a MutH nicking assay in the presence of full length MutL or MutL^LN40^ alone (Fig. 1C). As shown before, MutH shows no activity in isolation but is strongly activated in the presence of MutL and ATP or AMPPNP (8). The stronger activity of MutH in the presence of AMPPNP may be explained by the fact that this analogue cannot be hydrolysed and therefore retains more MutL molecules in the active, dimeric form, when compared to ATP. Next, when we use the MutL^LN40^ domain only (Fig. 1C, right panel), we see a similar MutL-ATP/AMPPNP-dependent activation but weaker than for the full length MutL protein. To determine if the reduced activation of MutH by MutL^LN40^ is due to a lack of a region located in MutL^LC20^ or due to a reduction of MutL^LN40^-dimers we performed two experiments. First, we added the isolated LC20 domain including the flexible linker (i.e. MutL residues 350-615) to a reaction containing MutL^LN40^ and MutH (Fig. 1D-E). Across all concentrations of MutL^LC20^ used, we observed no increased activity, even at 10 µM MutL^LC20^ dimer concentration. Next, we used size exclusion chromatography to follow the AMPPNP-induced dimerization of the LN40 domain at different time points (Fig. 1F). For the full length MutL protein, dimerization of the MutL^LN40^ domain results in compacting of the dimer, leading to a shift to the larger volumes in the size exclusion column (Supplemental Fig. S3). In contrast, the AMPPNP-induced dimerization of MutL^LN40^ creates a larger complex leading to a shift to the smaller volumes in the size exclusion column (Supplemental Fig. S3). For the full-length protein, an almost complete conversion to the more compact dimeric form is observed at 10 minutes which is identical to 85 minutes. In contrast, the isolated MutL^LN40^ domain shows only 7% dimers after 10 minutes, and ∼50% dimers after 85 minutes. Taken together, this suggests that the reduction of MutH activation by MutL^LN40^ is not due a missing part in MutL^LC20^ but due to a reduction in MutL^LN40^ dimers. Importantly, these results also indicate that only MutL^LN40^ is required for activation of MutH and that therefore our cryo-EM structure contains all the information needed to understand how MutL activates MutH.

### DNA binding in the MutH-MutL complex

In our structure of MutH-MutL-DNA, MutH is bound to a groove in between the two monomers of MutL^LN40^ (Fig. 1A and 2A) in agreement with earlier biochemical results (7, 8, 12). Interestingly, in a recent structure of MutL^LN40^ bound to a DNA substrate with a 5’ extended single stranded overhang (21), the same groove was occupied by the single stranded portion of the DNA substrate (Fig. 2B). This indicates that MutL^LN40^ cannot bind MutH and ssDNA at the same time, and hence, in the MutL-MutH complex, the DNA is only bound by MutH. Due to the binding of MutH, the four helices that cradle the groove between the two MutL^LN40^ domains are separated by 28 Å, which is 7 Å and 13 Å wider than the apo structure or DNA-bound structure of MutL^LN40^, respectively (Fig. 2C).

**Figure 2.**
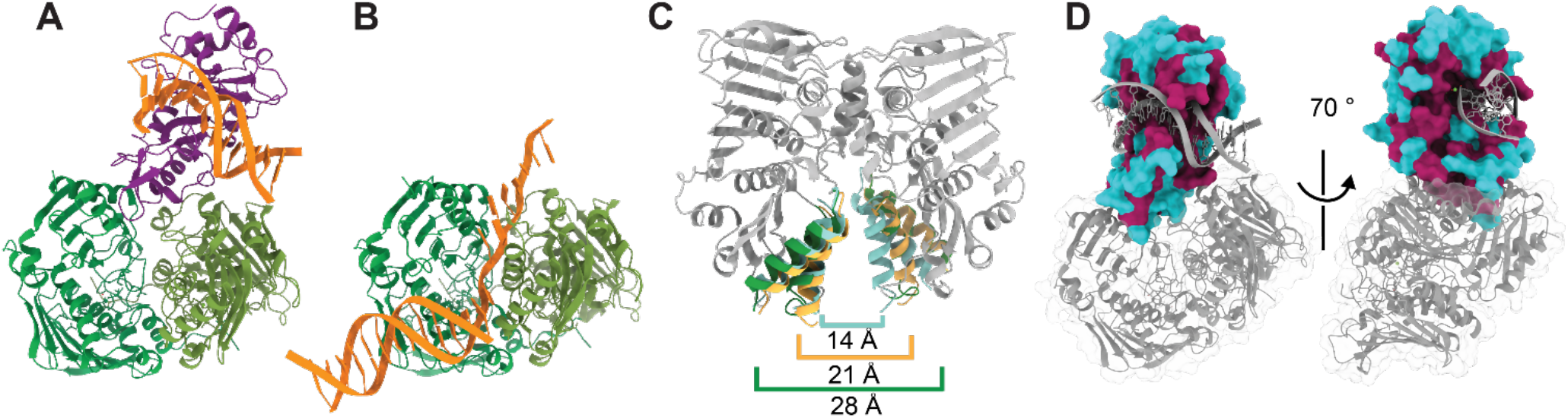
DNA binding in the MutL-MutH-DNA complex. **(A)** Structure of the MutL-MutH-DNA complex. MutL^LN40^ is coloured in light and dark green, MutH in purple, and the DNA in orange. **(B)** Structure of MutL^LN40^ bound to a 5′ extended single stranded DNA (PDB ID: 7P8V). In this structure, the single-stranded DNA overhang is located in the same groove as MutH in the MutL-MutH-DNA complex, indicting that MutL^LN40^ cannot bind MutH and DNA simultaneously. **(C)** Superposition of three MutL^LN40^ structures in different functional states: apo, ssDNA-bound, and MutH-bound. The changes in the groove between the two monomers are highlighted by helices 265–281 and 313–331. The width of the groove ranges from 14 Å in the ssDNA-bound structure (blue, PDB ID: 7P8V), to 21 Å in the apo structure (orange, PDB ID: 1B63), to 28 Å in the MutH-bound structure (green, this work. **(D)** Conservation analysis of MutH from *H. influenzae* and *E. coli* mapped onto the *E. coli* MutH structure. Highly conserved residues are shown in magenta, and lesser conserved residues in blue. The strong conservation of the DNA-binding site supports the use of the available *H. influenzae* MutH-DNA structure (PDB ID: 2AOR) as a structural reference for comparison with our *E. coli* MutL-bound MutH complex.

For our structure we used a 14-base pair DNA substrate, identical to the substrate that was used in the MutH activity assays (Fig. 1C-D). The DNA is bound in a manner similar to that of the *Haemophilus influenzae* MutH-DNA complex in absence of MutL (Lee Yang 2002), with the DNA slightly bent and grasped between three loops (Fig. 2A and 2D. The sequences of *H. influenzae* and *E. coli* MutH are 80% similar but show almost 100% conservation in the DNA binding region and MutL-interacting region (Fig. 2D, Supplementary Fig. S1) and can therefore be used to compare changes between the MutL-free and MutL-bound MutH-DNA complex.

### The pre-incision state of MutH

The active site of MutH is well resolved in the cryo-EM map, with the magnesium ions and coordinated water atoms clearly visible (Fig. 3A). Compared to the MutL-free structure of MutH-DNA, the DNA has moved 1.0 Å closer to the MutH active site (Fig. 3B). This movement of the DNA is stabilized by histidine 112 that in its new position hydrogen bonds to the oxygen of the DNA backbone located immediately 5’ to the cut site in the DNA. In the new position, lysine 116 positions a water molecule perfectly in line with the scissile bond 5’ of the GATC sequence that is to be cut. Lysine 116 is strictly conserved in MutH homologs and its mutation renders MutH inactive (22). However, catalysis does not take place as glutamate 56 that contributes to the coordination of metal B, is mutated to an alanine (Fig. 3B, Supplemental Fig. S1).

**Figure 3.**
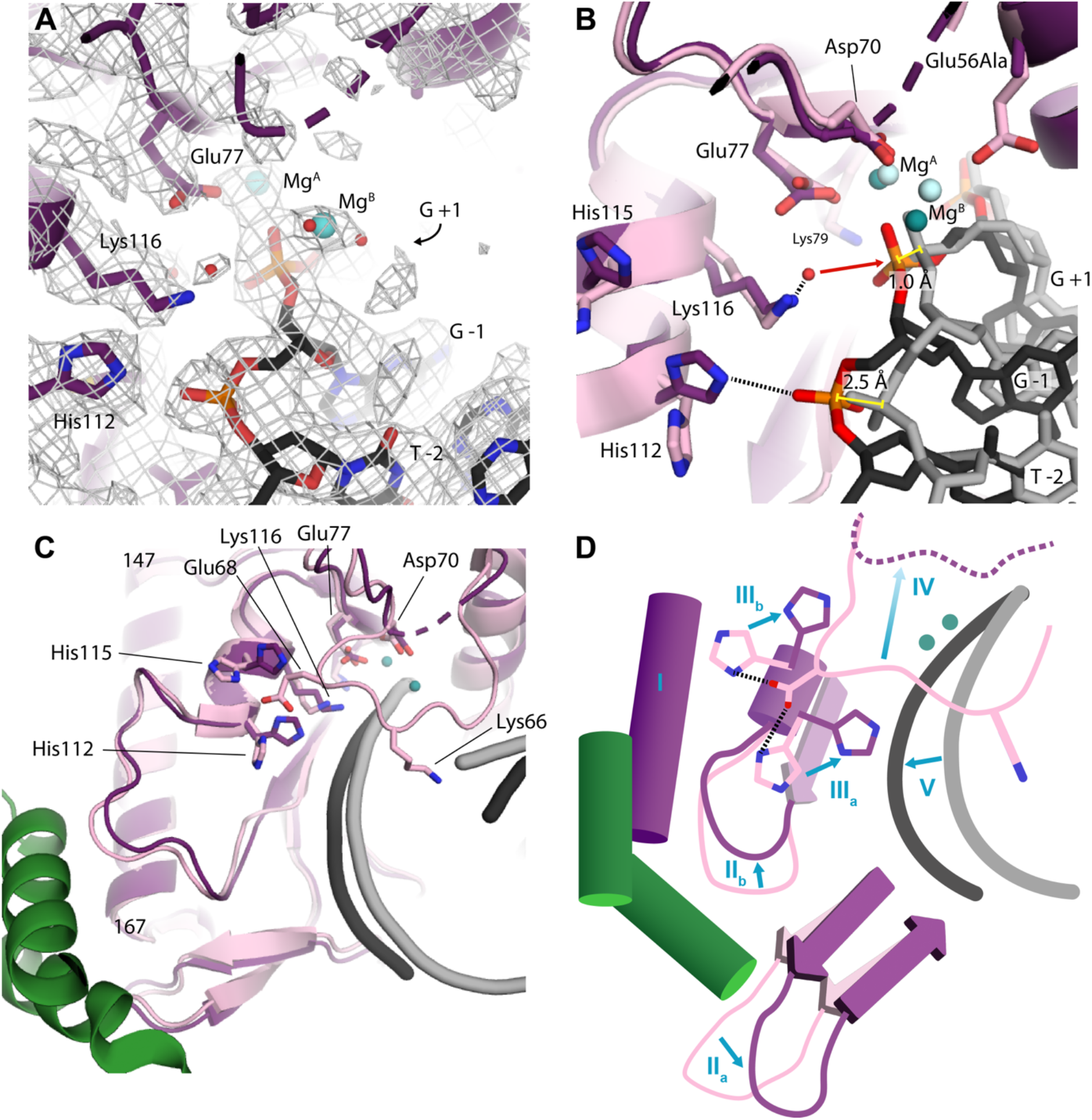
Activation of MutH by MutL. **(A)** Active site configuration of MutL-bound MutH. Cryo-EM map is represented in grey mesh, magnesium ions as blue spheres and water molecules as red spheres. Key residues are shown in sticks. Numbering of the bases are relative to the scissile bond. **(B)** Comparison of the MutL-bound structure (in dark colours) and MutL-free structure (in light colours) of MutH-DNA. Key residues and the DNA are shown in sticks. Hydrogen bonds are shown as black dashed lines, the possible angle of attack of the water molecule to the scissile phosphate is shown by a red arrow. Yellow lines mark the movement of the -1 and +1 phosphates from the MutL-free to the MutL-bound form. **(C)** Structural rearrangements in MutH upon MutL binding. Super positioning of the MutL-interacting helix of MutH (spanning residues 147-167) reveals the conformational changes that connect MutL binding to the movement of the DNA into the MutH active site. **(D)** Schematic representation of the allosteric pathway that couples MutL-binding to MutH active site, 25 Å away. (I) MutL binds MutH at a helix spanning residues 147-167. (II) The binding of MutL causes the movement of a β-hairpin downwards (II_a_) and a loop upwards (II_b_). (III) The loop movement coincides with the rotation of two histidines (His112 and His115 - III_a_ and III_b_) that form hydrogen bonds to a glutamate (Glu68), which is part of a loop that interacts with the minor groove of the DNA. (IV) Upon release of the hydrogen bond to Glu68, the DNA binding loop moves away from the DNA and becomes disordered. (V) As a result, the DNA backbone can move closer into the active site (marked by two green spheres), stabilized by a hydrogen bond to histidine 112.

### Activation of MutH by MutL

MutL^LN40^ binds to MutH via a helix spanning residues 147-167, located ∼25 Å away from the active site (Fig. 3C). No direct contacts are made to the active site of MutH, implying that the activation could be achieved through allosteric regulation. To reveal how MutL binding induces changes in MutH we superimposed the MutL-binding helix of MutH (residues 147-167) of the MutL-free and MutL-bound structures (Fig. 3C). This reveals that MutL acts on MutH through a ‘push-and-pull’ mechanism, depicted schematically in Fig. 3D. A β-hairpin in MutH (residues 192-207, labelled ‘IIa’) is pulled down by MutL, while an adjacent loop (residues 100-110, labelled ‘IIb’) is pushed up. The upwards motion of this loop coincides with the flipping of histidines 112 and 115 towards the DNA (labelled ‘IIIa and IIIb’). In this new position, the histidines no longer hydrogen bond to glutamate 68, which is part of a loop (residues 62-69) that positions lysine 66 in the minor groove of the DNA in the MutL-free form and appears to keep the DNA in place (Fig. 3C). In the MutL-bound form of MutH-DNA the glutamate 68 is no longer held in place and consequently the loop and lysine 66 have moved out of the DNA minor groove and become flexible as evidenced by its absence in the cryo-EM map (labelled “IV”). With the loop out of the way, the DNA is free to move deeper into the active site of MutH (labelled ‘V’), where it is stabilized by histidine 112 that has flipped ∼180° from its position in the MutL-free form. This way, the binding of MutL ∼25 Å away from the cut site in the DNA, activates MutH through an allosteric network that releases an inhibitory loop from the minor groove of the DNA, while pulling the DNA closer into the active site through the flipping of histidine 112.

### A conserved mechanism of activation in Type II restriction endonucleases

The activation of MutH by MutL bears similarities to the activation mechanism of Sau3AI, a GATC-specific type II restriction endonuclease. The N- and C-terminal domains of Sau3AI are both homologous to MutH. In Sau3AI, the N-terminal domain acts as the MutH-like catalytic nuclease module, while the C-terminal domain acts as a non-catalytic effector. Structural analysis of Sau3AI indicates that, in the absence of a substrate, the catalytic cleft of the N-terminal domain is obstructed by the C-terminal domain, resulting in an autoinhibited state. It is then proposed that binding of a first GATC site to the C-terminal effector domain triggers a domain reorientation that opens the catalytic cleft of the N-terminal domain and permits cleavage at a second GATC site (23). Therefore, like MutH, it appears that Sau3AI does not achieve full catalytic competence through substrate binding alone, but rather through distal allosteric input that relieves inhibition and enables DNA cleavage.

Our MutL-activated MutH structure suggests the same general principle but implemented in trans rather than in cis. In our complex, MutL binds MutH approximately 25 Å from the active site. Through the push-and-pull rearrangement described above, MutL promotes the release of an inhibitory loop from the DNA minor groove and the movement of the DNA towards the active site. Therefore, Sau3AI may represent an evolutionary reworking of the same regulatory logic: in Sau3AI, a built-in MutH-like effector domain senses the DNA and opens the catalytic cleft. In contrast, in MutH, the activating signal is supplied by the ATP-bound MutL dimer.

## Discussion

Mismatch repair relies on a series of carefully coordinated interactions among MutS, MutL, and MutH that ensure that after mismatch recognition incision occurs at the newly synthesized strand (Fig. 4). While the early stages of mismatch recognition and MutS– MutL signaling are structurally well defined (21, 24–27), how MutL subsequently activates MutH remains poorly understood. Our MutL–MutH structure now provides a detailed understanding how conformational rearrangements in MutH are coupled to its binding by MutL, bridging the communication gap between mismatch detection and strand incision.

**Figure 4.**
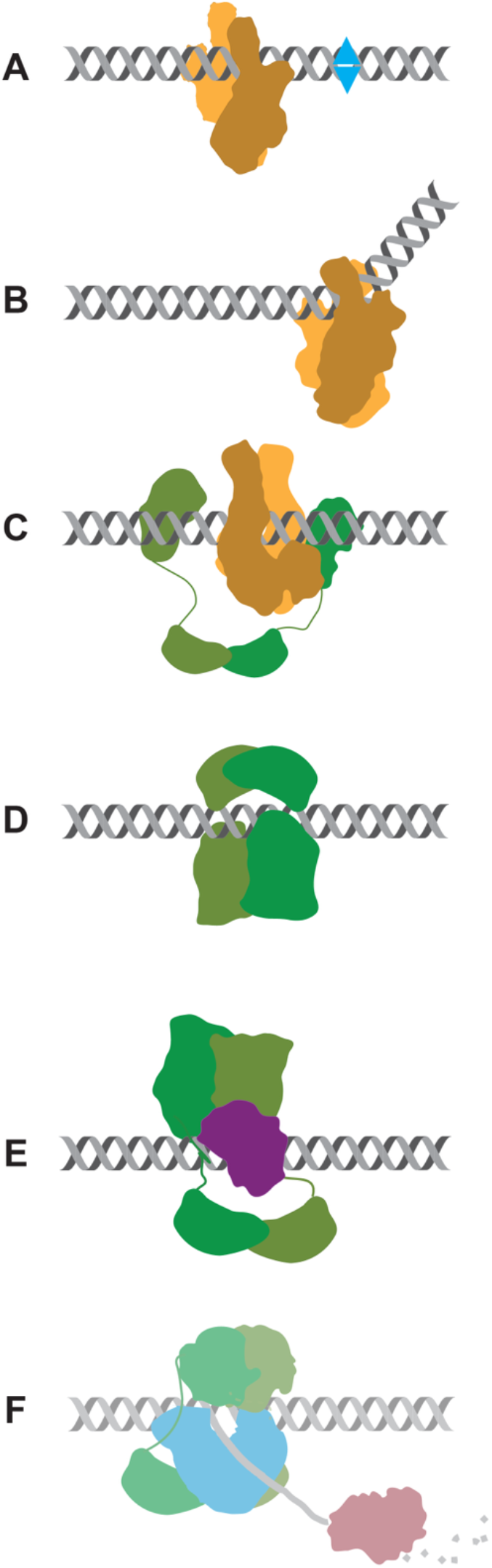
The sequential steps of the DNA mismatch repair cascade. **(A)** MutS (coloured in yellow-orange) scans the DNA searching for a mismatched base pair (coloured in blue). **(B)** After mismatch recognition, MutS undergoes an ATP-induced conformational change that **(C)** recruits MutL (coloured in green) to the DNA. **(D)** In turn, MutL also undergoes an ATP-induced conformational change that results in a closed MutL dimer that can scan the DNA in search of MutH bound to a hemi-methylated GATC-site. **(E)** MutL will then activate MutH (coloured in purple) to nick the newly synthesized strand, after which **(F)** MutL will guide the helicase UvrD (coloured in blue) to unwind the DNA and expose the newly synthesized strand to an exonuclease (coloured in brown). The structural features of the MutL-UvrD interaction remain to be determined.

First, MutS scans the DNA (Fig. 4A) and after mismatch recognition (Fig. 4B) undergoes an ATP-dependent conformational change that enables it to recruit MutL and load it onto DNA (Fig. 4C). MutL then also undergoes an ATP-dependent conformational change into a clamp-like state (Fig. 4D) that diffuses along the DNA (28, 29). This ATP-bound dimeric form of MutL represents the active signaling state that engages with MutH. At this stage, one could envision two possible order events: (i) MutL and MutH form a complex on DNA that together search for a hemi-methylated GATC site, or (ii) MutL travels on the DNA alone in search for a MutH that is pre-bound to a hemi-methylated GATC site. The affinity between MutH and MutL however appears to be too low, as even at 22 µM there is limited interaction between the two molecules as judged by size exclusion chromatography (Supplemental Fig. S4). In contrast, the affinity of MutH for a hemi-methylated GATC site is 0.9 µM (7), suggesting that MutL travels alone on the DNA in search of MutH bound to a hemi-methylated GATC site. Finally, after strand incision, the helicase UvrD is recruited to the nick and together with one of four exonucleases RecJ, Exo I, Exo VII, or ExoX will resect the newly synthesized strand between the nick and the mismatch (30, 31). However, the structural features of the MutL-UvrD interactions remains to be determined.

## SUPPLEMENTARY INFORMATION

Supplementary data is available at NAR online.

## ACKNOWLEDGEMENTS

We thank The Netherlands Center for Electron Nanoscopy (NeCEN) for cryo-EM data collection. We thank Willem Noteborn for suggestions on cryo-EM data processing, and Rafael Fernandez Leiro and Luuk Loeff for helpful discussions.

## Author contributions

E.A.U. and M.H.L. designed the project. E.A.U. preformed experiments, collected and processed cryo-EM data. E.A.U. and M.H.L. wrote the manuscript.

## DATA AVAILABITILY

The cryo-EM density map of the MutL-MutH-DNA complex has been deposited in the Electron Microscopy Data Bank (EMDB) under accession code EMD-57524. The atomic coordinates of the MutL-MutH-DNA complex has been deposited in the Protein Data Bank (PDB) under the accession code 30AX. .

