## Supplemental for "Cryo-EM structure of MutL-activated MutH"

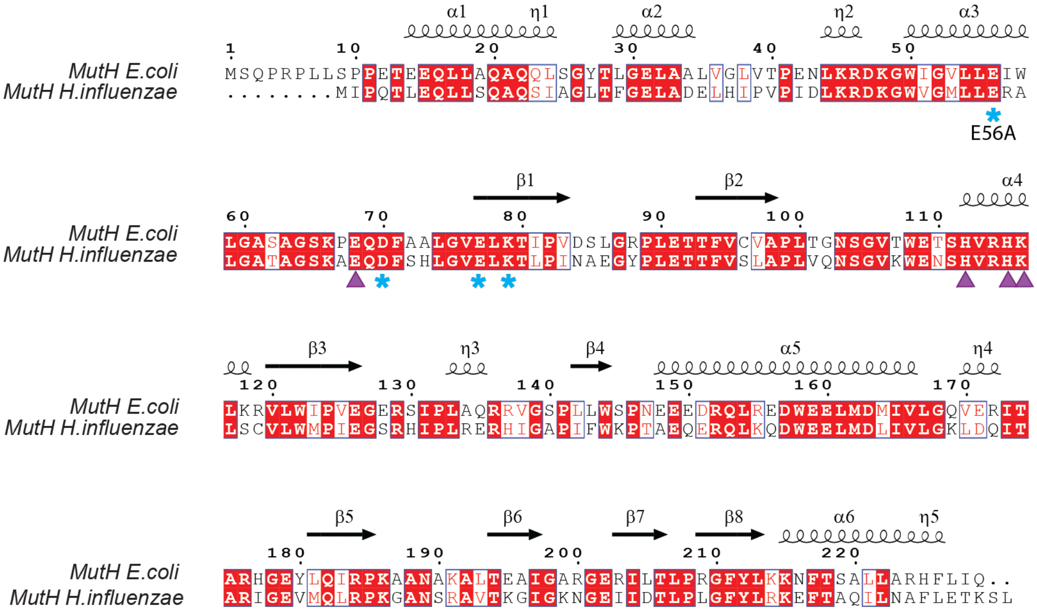


**Figure S1. Sequence conservation between *E. coli* and *H. influenzae* MutH.** Sequence alignment of MutH from *Escherichia coli* and *Haemophilus influenzae*. Conserved residues are highlighted in red background, similar residues in red letter and white background. Active-site residues are indicated by blue stars. Residues discussed in this study are marked with purple triangles. (Related to main Figure 2D).

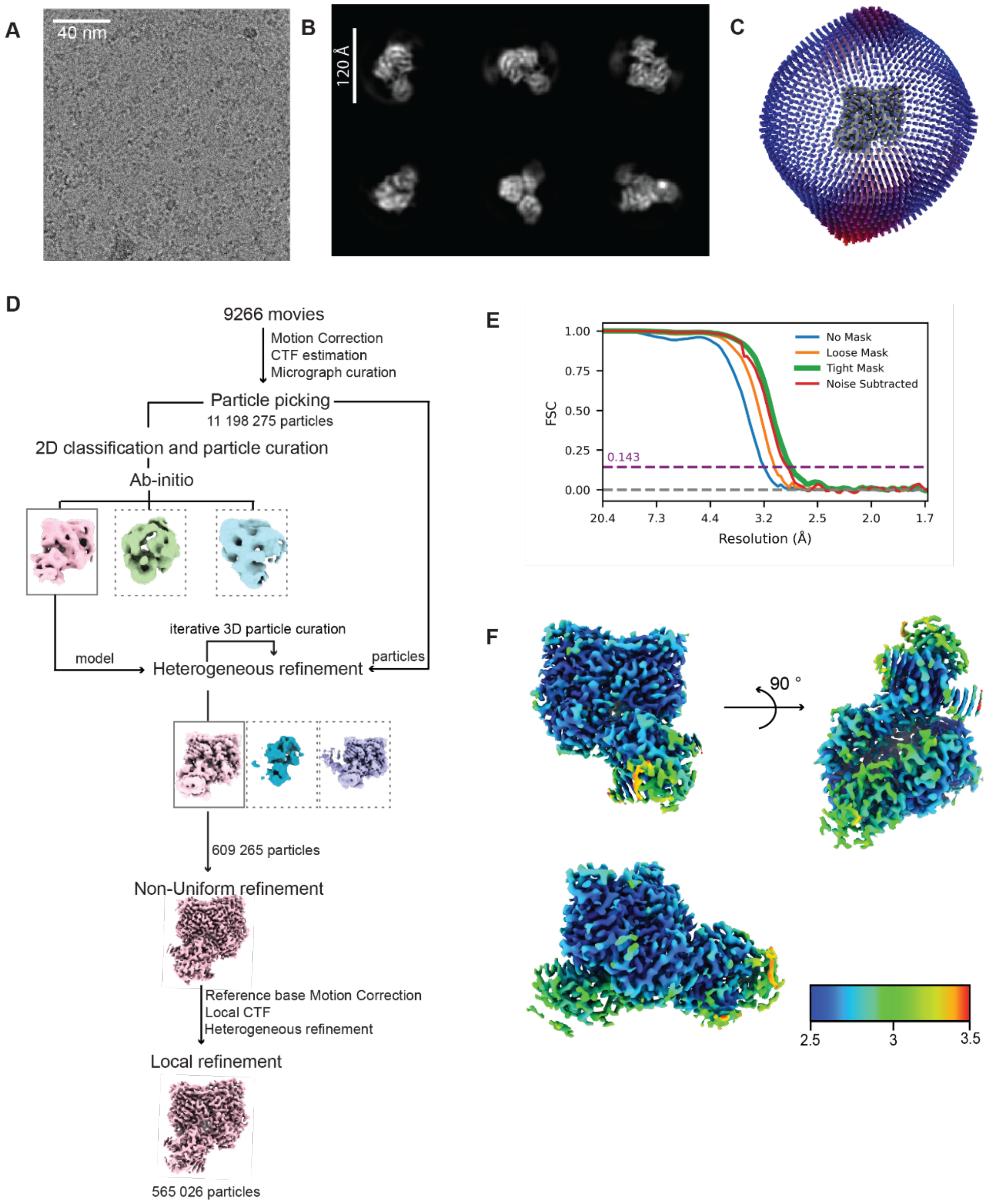


**Figure S2. Cryo-EM data processing overview of the MutL–MutH-DNA complex. (A)** Representative cryo-EM micrograph from the dataset. **(B)** Representative 2D class averages of the MutL–MutH-DNA complex. **(C)** Angular distribution of particles contributing to the final reconstruction. **(D)** Schematic representation of main data processing workflow. Volumes outlined with a continuous line indicate the reconstruction that was selected for further refinement and yielded the final map. **(E)** Fourier shell correlation (FSC) curve between half-maps from the final refinement used to estimate the final resolution. **(F)** Final refined map colored by local resolution.

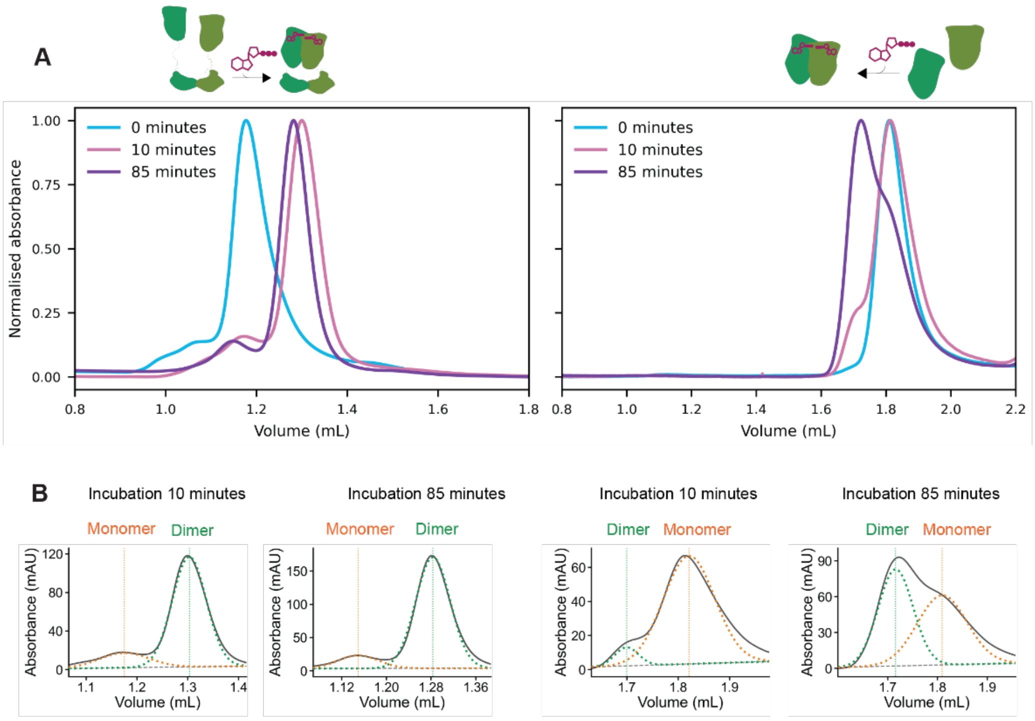


**Figure S3. Dimerisation of MutL$^{FL}$ and MutL$^{LN40}$. (A)** Size-exclusion chromatography profiles of full-length MutL incubated without nucleotide or with AMPPNP for 10 or 85 minutes prior to injection. In the presence of AMPPNP, the elution peak shifts toward a higher elution volume (corresponding to a more compact conformation). **(B)** Size-exclusion profiles of the isolated LN40 domain incubated without nucleotide or with AMPPNP for 10 or 85 minutes. A small shoulder at the retention volume of the dimer species is observed after 10 minutes of incubation that increases after 85 minutes incubation. **(C-D)** Quantification of monomeric and dimeric MutL population through fitting of a sum-of-two-Gaussians to the size exclusion profiles shown in panels A and B. (Related to main Figure 1F)

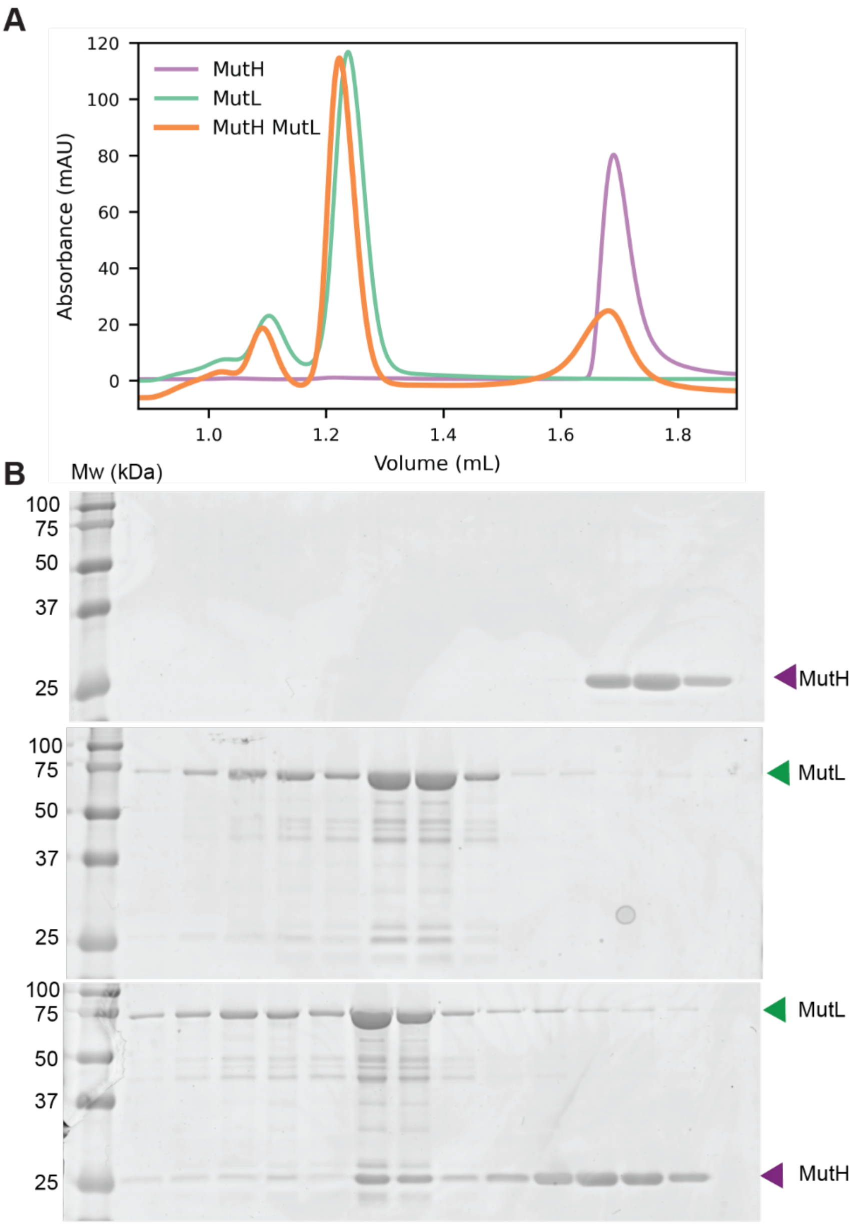


**Figure S4. MutL and MutH binding in absence of DNA. (A)** Size-exclusion chromatography profiles of 22 μM MutH (purple), 22 μM MutL dimer (green) and 22 μM MutH-MutL$_{dimer}$ combined. All reaction were performed in presence of 3 mM AMPPNP. **(B)** SDS-PAGE analysis of fractions from the size exclusion chromatography runs. Note the partial shift of MutH molecules to the retention volume of MutL, indicative of a weak interaction at 22 μM.

| | MutL-MutH-DNA |
|---|---|
| **Data collection** | |
| Microscope | Titan Krios |
| Voltage (kV) | 300 |
| Magnification | 105,000x |
| Detector - GIF | Gatan K3 - Bioquantum |
| Data collection software | EPU |
| Electron exposure ($e^{-}/Å^2$) | 60 |
| Defocus range (μm) | 0.8 - 2 |
| Pixel size (Å) | 0.836 |
| **Data processing** | |
| Software | CryoSPARC v4.7.0 |
| Number of micrographs | 9266 |
| Final number of particles | 565 026 |
| Symmetry imposed | C1 |
| Map resolution (Å) | 2.8 |
| FSC threshold | 0.143 |
| **Model Refinement** | |
| Software | Phenix 1.18 |
| Map correlation coefficient | 0.85 |
| Model composition | |
| Number of chains | 5 |
| Non-hydrogen atoms | 7481 |
| Protein residues | 877 |
| Ligands | 7 |
| **Validation** | |
| R.M.S. deviations | |
| Bond lengths (Å) | 0.013 |
| Bond angles (°) | 1.998 |
| Ramachandran plot | |
| Favored (%) | 96 |
| Allowed (%) | 4 |
| Disallowed (%) | 0 |
| Rotamer outliers (%) | 0 |
| **Data availability** | |
| EMDB entry | EMD-57524 |
| PDB entry | 30AX |

Supplemental Table 1. Cryo-EM data collection and refinement
